# Robust and Interpretable PAM50 Reclassification Exhibits Survival Advantage for Myoepithelial and Immune Phenotypes

**DOI:** 10.1101/480723

**Authors:** James C. Mathews, Saad Nadeem, Arnold J. Levine, Maryam Pouryahya, Joseph O. Deasy, Allen Tannenbaum

**Author notes:** **Correspondence:** James C. Mathews.

## Abstract

We introduce a classification of breast tumors into 7 classes which are more clearly defined by interpretable mRNA signatures along the PAM50 gene set than the 5 traditional PAM50 intrinsic subtypes. Each intrinsic subtype is partially concordant with one of our classes, and the 2 additional classes correspond to division of the classes concordant with the Luminal B and the Normal intrinsic subtypes along expression of the Her2 gene group. Our Normal class shows similarity with the myoepithelial mammary cell phenotype, including TP63 expression (specificity: 80.8% and sensitivity: 82.8%), and exhibits the best overall survival (89.6% at 5 years). Though Luminal A tumors are traditionally considered the least aggressive, our analysis shows that only the Luminal A tumors which are now classified as myoepithelial have this phenotype, while tumors in our luminal class (concordant with Luminal A) may be more aggressive than previously thought. We also find that patients with Basal tumors surviving to 48 months exhibit favorable survival rates when certain markers for B-lymphocytes are present and poor survival rates when they are absent, which is consistent with recent findings.

## 1 Introduction

Multiparametric genetic tests such as the PAM50/Prosigna Risk of Recurrence (ROR) for breast cancer prognostication are becoming commonplace [1,2]. However, due to limited accuracy and poor concordance with biological phenotypes, their clinical utility is still under investigation [3]. In this paper we address these issues in the context of one of the most prevalent assays, the PAM50 ROR, which is mainly driven by an intrinsic subtype classification along a 50-gene mRNA expression profile. We reclassify these profiles using topological data analysis, incorporating prior knowledge of biological phenotype (basal/luminal stratification). Unlike the 5 traditional PAM50 intrinsic subtypes, our 7 classes are accurately defined by clear patterns of activation and inactivation of gene groups directly interpretable in terms of specific normal mammary cell types: **basal, luminal/ER, myoepithelial**, and **Her2-related** gene groups.

The basal/luminal terminology refers to mammary cell differentiation from basal-epithelial cells near the basement membrane to the more differentiated luminal-epithelial cells near the lumen or ducts. It was the basis for the systematic molecular classification of breast cancer initiated by Perou *et al*. [4]. Myoepithelial refers to a mammary cell type playing a key role in breast duct secretion [5,6]. Overexpression of Her2 (ERBB2) and a group of related genes marks the Her2+ cohort well-known since the 1990s for highly favorable response to the drug trastuzumab (herceptin). Figure 1 summarizes the history of the molecular classification and our contribution. Table 1 lists the new classes.

**Figure 1:**
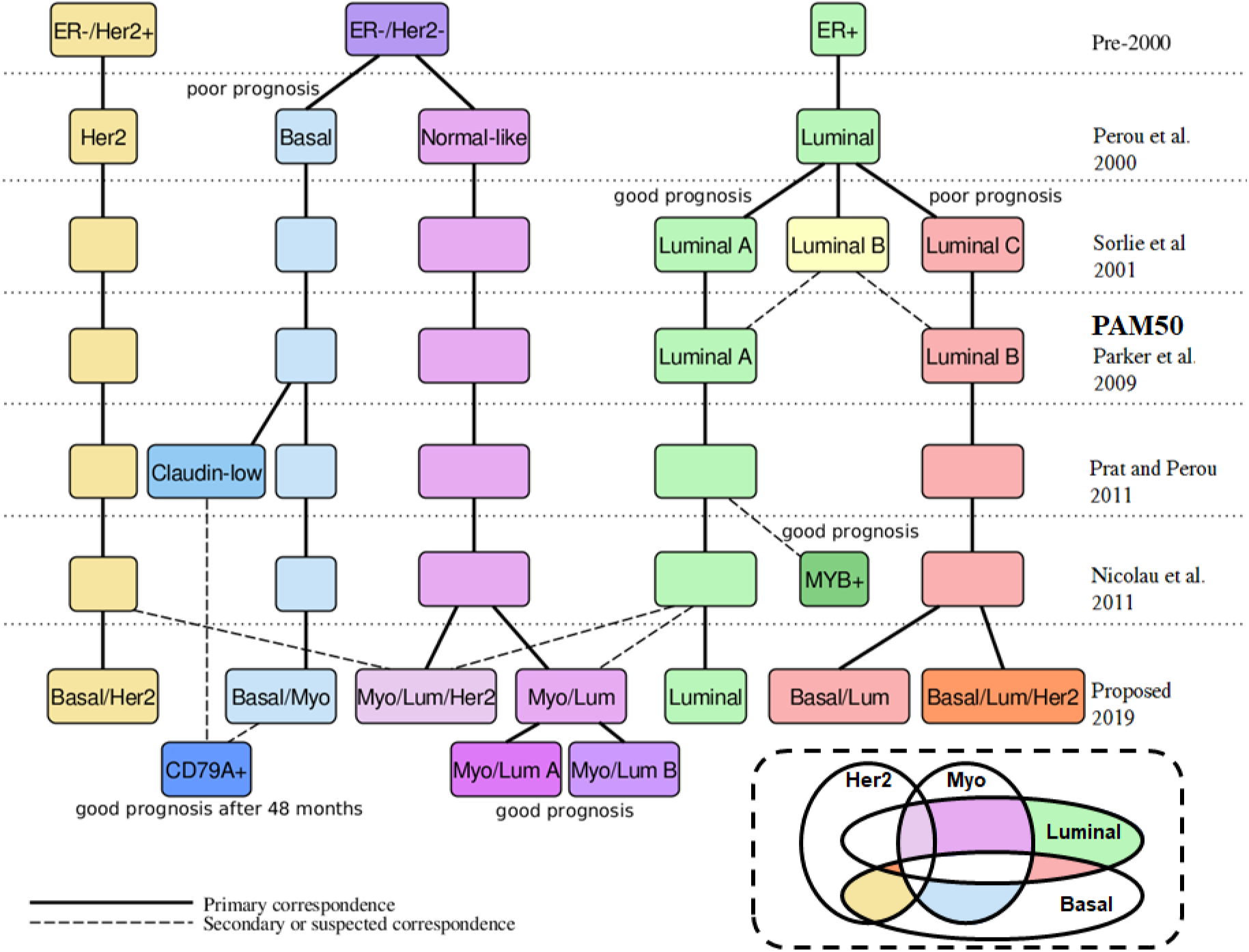
History of the molecular classification of breast cancer. Names are shown at the chronological level at which they were introduced. The Her2+ breast tumors were already well-known in the 1990s for highly favorable response to the drug trastuzumab (herceptin), which was approved by the FDA for metastatic Her2+ breast cancer in 1998. The hierarchical clustering of Perou *et al*. [4] used genes whose expression differentiates between samples from different tumors better than between samples from the same tumor, finding 4 main classes: ERBB2+ (or Her2+), Basal, Luminal, and Normal-breast-like. Sorlie *et al*. [7] explicitly incorporated clinically relevant outcome data such as overall survival, uncovering three Luminal subtypes, Luminal A, B, and C. Luminal A has higher overall survival than Luminal B and Luminal B has higher overall survival than Luminal C. Later investigators found only two Luminal subtypes to be sufficiently robust. Parker *et al*. [8] introduced the 50 gene set that became known as the PAM50 (Prediction Analysis of Microarray) and introduced a straightforward centroid-based classifier for breast tumor RNA expression patterns along the PAM50 with 5 classes: Basal, Her2, Luminal A, Luminal B, and Normal. The authors used this classification as a key component in the model that became the Prosigna predictor of Risk Of Relapse (ROR). Prat and Perou [9] introduced the Claudin-low subtype carved largely out of the Basal group. The authors find that the Claudin-low subtype has poor prognosis compared to Luminal A, but no worse than the other subtypes. The Topological Data Analysis of Nicolau *et al*. [10] confirmed the distinction between more luminal, more basal, and more normal-like subtypes along branches of a graph structure modeling the distribution of breast tumor samples. They found a subgroup of patients exhibiting a very high survival rate, largely characterized by expression of MYB. **Our proposed classification uses the method of Nicolau *et al*. [10] and incorporates gene sets and priors (e.g. the basal-to-luminal stratification) known to be relevant to breast cancer biology**. (Below right) Our proposed system with 7 classes defined by 4 elementary phenotypes (see also Figures 3 and 2).

**Table 1:**
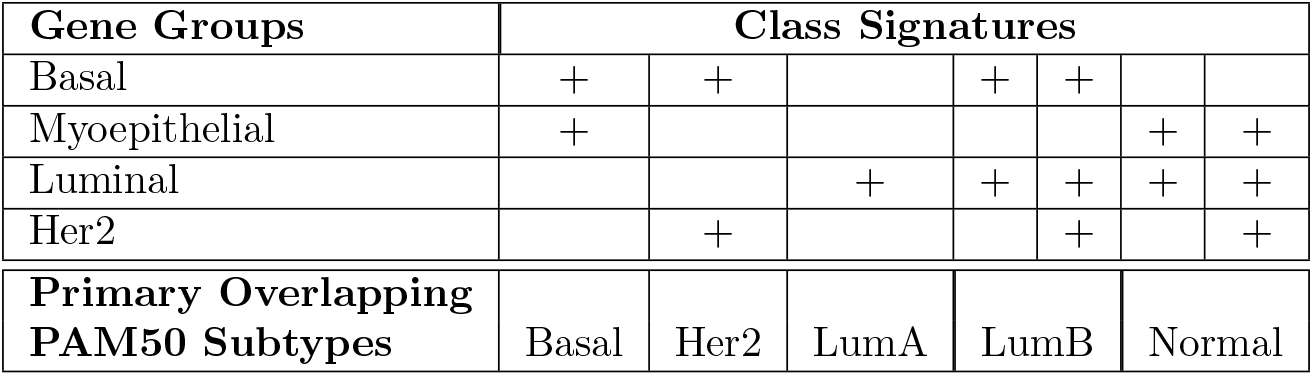
Reclassification of PAM50 subtypes of breast tumors. *The genes in each gene group are shown in Figure 2*.

## 2 Results

### Clearly-defined 50-gene signatures (Figure 2)

The signature classes we defined show partial concordance with the PAM50 subtypes, with a Normalized Mutual Information (NMI) of 0.19 (29.1 times the maximum NMI found in 10000 random permutation bootstrapping trials). However, our classes show tighter clustering along the 50-gene profile: the k-mean for the PAM50 subtypes is 87.9% of the total variance, and for our classification is only 82.7% (both using the L1 norm). To assess the quality of the signatures themselves, we consider the *average silhouette width* [11] of each class. The silhouette width is the average distance *a(i)* between a sample *i* and the cluster to which it belongs subtracted from the smallest average distance *b(i)* between *i* and the other clusters, normalized by *max*(*a*(*i*), *b*(*i*)). The average silhouette width over a given cluster (abbreviated SW) measures the tightness of the cluster with respect to the clustering scheme, with larger SW (closer to 1) indicating a good cluster and smaller SW (closer to −1) indicating a poor cluster.

**Figure 2:**
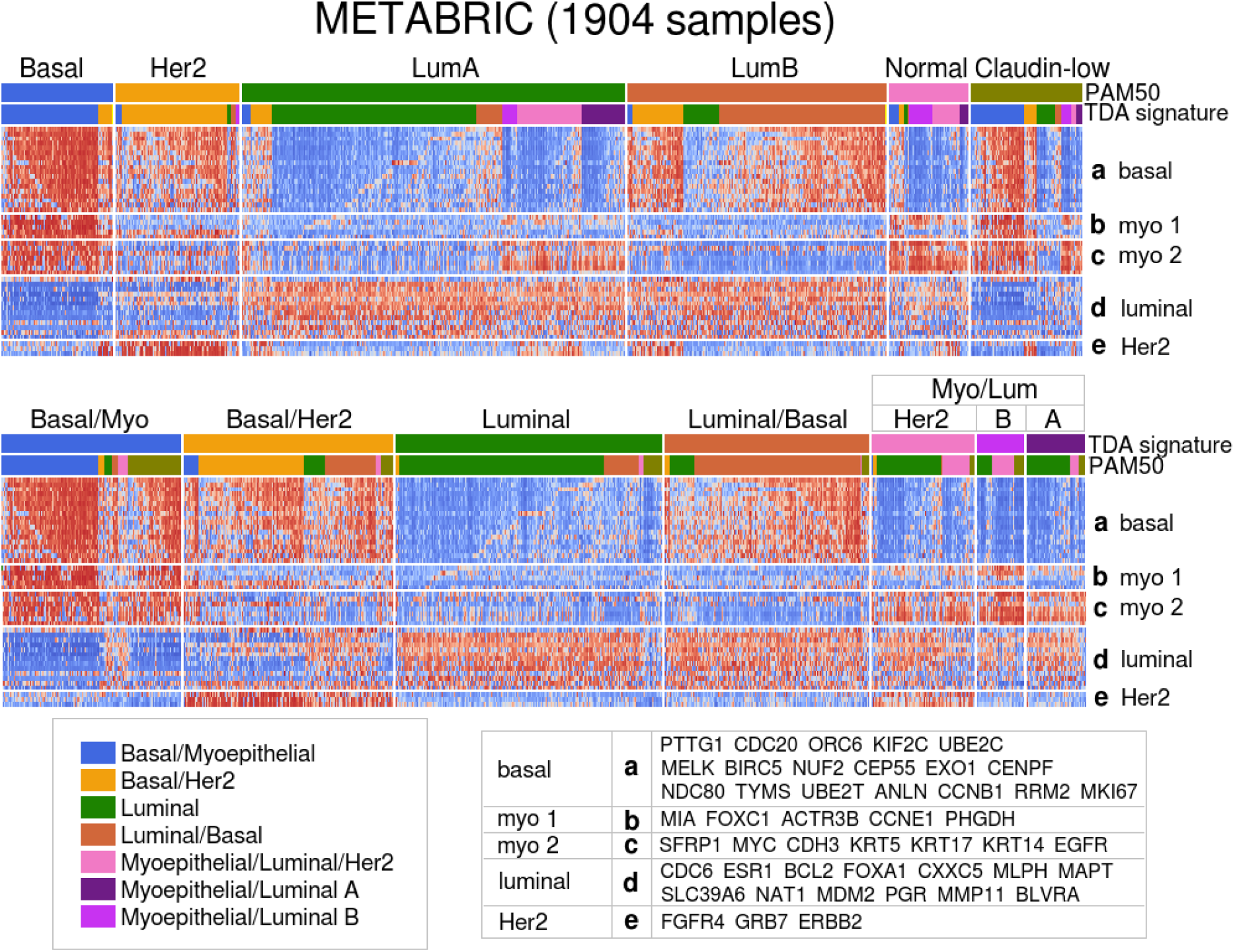
The RNA expression heatmap of the 1904 METABRIC breast tumor samples. (Above) Organized first by PAM50 subtype and then by the TDA signatures classes assigned by the Mapper-derived classifier along the PAM50 gene set (BAG1, MYBL2, GPR160, and TMEM45B omitted due to missing values). (Below) Organized first by TDA signature class then by PAM50 subtype.

**Figure 3:**
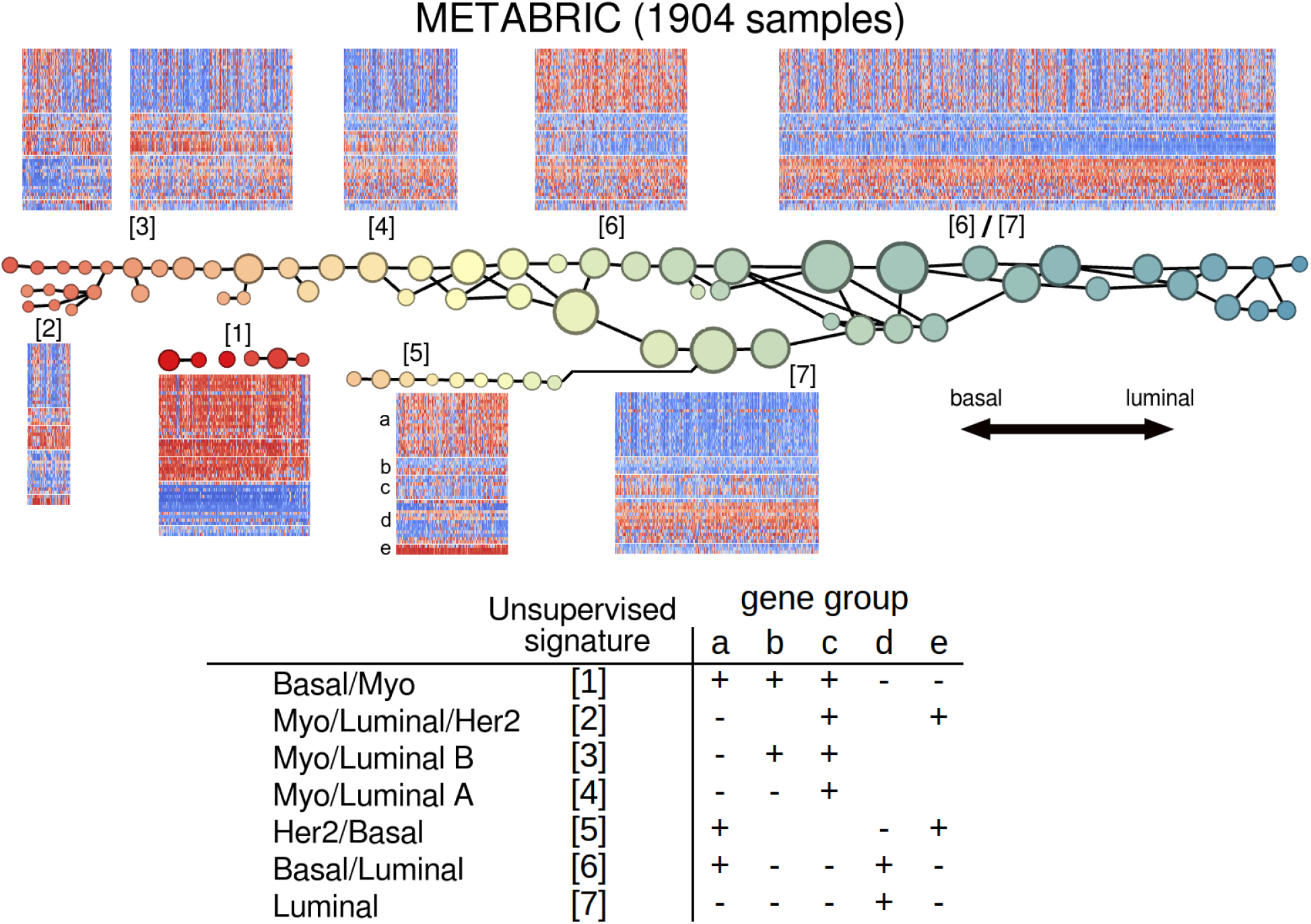
(Above) The Mapper analysis of the 1904 METABRIC breast tumor samples, along the PAM50 gene set, using the basal-luminal score as filter function. The circular nodes represent clusters in the strata or bins defined by the filter function at the chosen level of resolution. For example, there are 3 clusters in the stratum shown in yellow; two clusters shown higher and labelled with unsupervised signature number [4], and one cluster shown lower labelled with unsupervised signature number [5]. All 3 have the same basal-luminal score range, indicated by color. (Below) The salient signatures recorded. These signatures differ slightly, in two ways, from the 7 classes we finally propose as in Figure 1. First, for the sake of simplicity we merge the 2 myoepithelial-related gene groups (b) and (c) into a single gene group, consequently merging Myo/Luminal A and Myo/Luminal B into Myo/Luminal. Second, on account of the salient signatures observed in the heatmaps in Figure 2, we split Her2/Basal [5] into Her2/Basal and Luminal/Basal/Her2. Where blanks appear, the corresponding gene group (a)-(e) is neither uniformly positively nor uniformly negatively expressed.

Our Luminal class SW = 0.151 is greater than the PAM50 Luminal A SW by 0.107; Luminal/Basal SW = 0.131 is greater than the PAM50 Luminal B SW by 0.112; Myo/Luminal SW = 0.0422 is greater than the PAM50 Normal SW by 0.0432 (silhouette widths range from −1 to 1). The SWs of our Her2 and Basal/Myo SWs are very close to the SW of the PAM50 Her2 and Basal subtypes.

As shown in Figure 2, the main example of a clear new signature is the heterogeneous expression of the myoepithelial gene group in the PAM50 Luminal A subtype, resolved by division into Luminal and Myo/Luminal classes. One exception is that the Basal/Her2 class binds together the PAM50 Her2 with several PAM50 Luminal B samples. However, the Luminal B here clearly differ from the Her2 by the presence of Luminal markers, so to address this we divide this class into Basal/Her2 and Basal/Her2/Luminal. Also, the two myoepithelial gene groups are small and closely related, so we merge them together into a single myoepithelial group and accordingly merge the classes denoted Myo/Luminal A and Myo/Luminal B. The 7 resulting signatures are shown in Table 1.

### Myo/Luminal class with good survival (Figures 4, 5, and 6)

The Kaplan-Meier survival analysis of the new classes is shown in Figure 4 for both 1904 METABRIC and 1082 TCGA samples. The plots show that the Myo/Luminal class exhibits the greatest survival rate, even greater than PAM50 Luminal A (the log-rank test for statistically significant difference between Normal and Myo/Luminal survival curves yields p = 0.003). Many of the Myo/Luminal tumors are designated PAM50 Luminal A, and since the Luminal A subtype is already the one with the best prognosis in the PAM50 scheme, we conclude that the Myo/Luminal class preferentially selects from Luminal A subtype the patients with especially good prognosis even among Luminal A.

**Figure 4:**
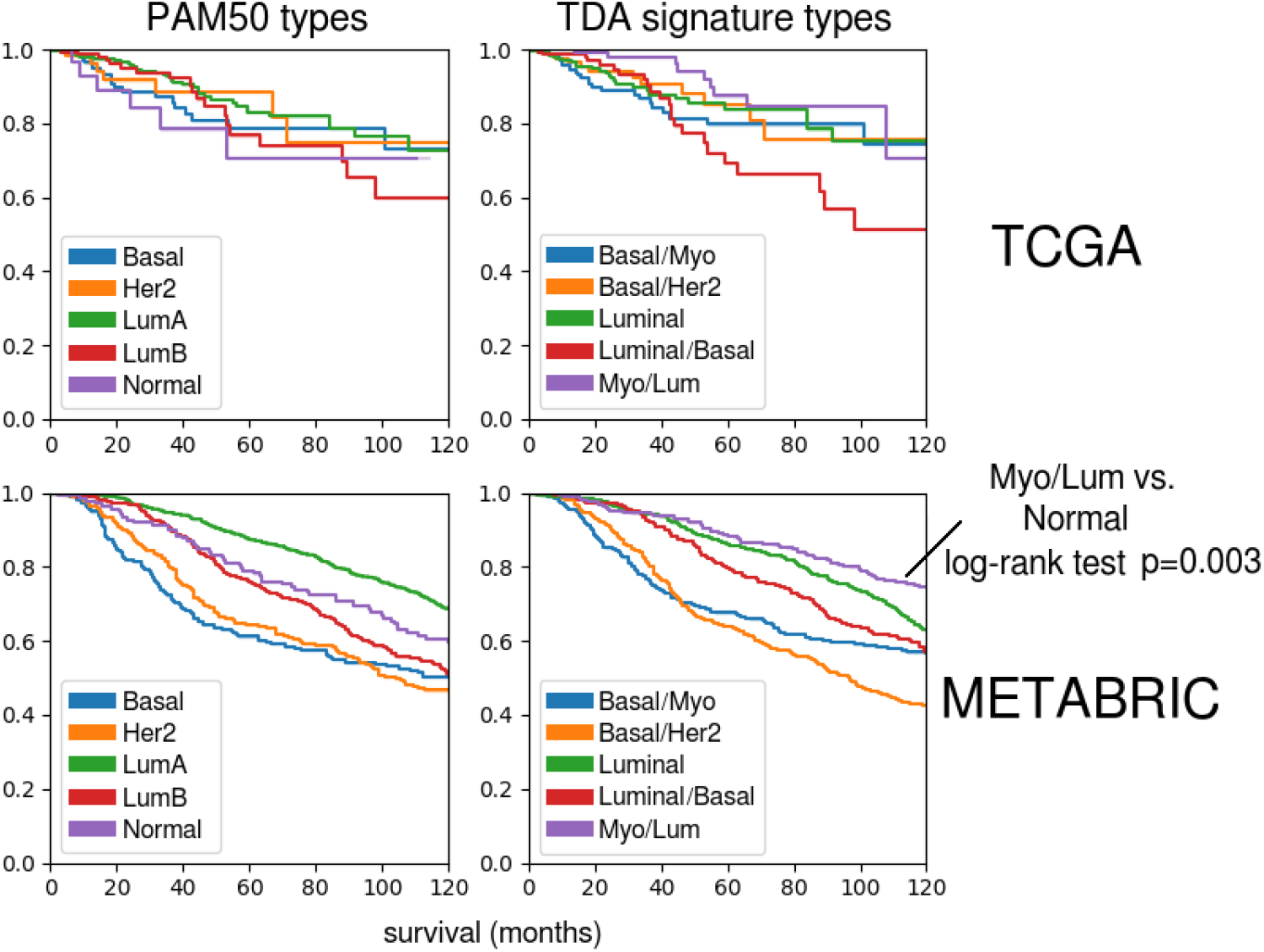
Kaplan-Meier survival analysis of the subgroups of the TCGA and METABRIC cohorts, respectively, defined by PAM50 subtypes and the major corresponding TDA signature classes, respectively. The Myo/Luminal class has the highest survival rate, statistically significantly greater than the primary corresponding PAM50 subtype, the Normal subtype. In the TCGA dataset, the log-rank test for PAM50 Normal versus Myo/Luminal yields p=0.023, while in the METABRIC dataset (with approximately twice as many samples) the test yields p=0.003.

**Figure 5:**
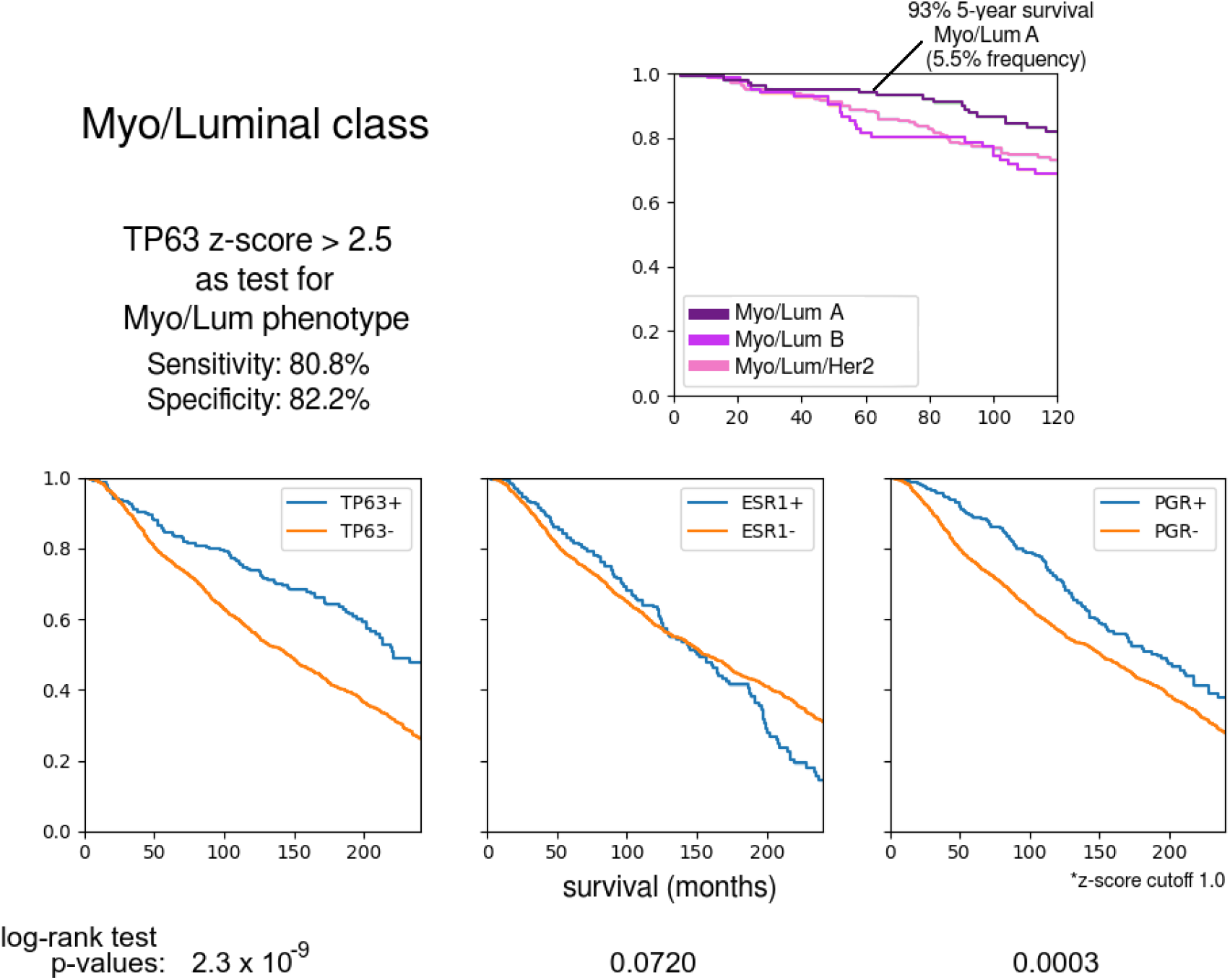
The stratification of the Myo/Luminal class by 3 TDA signature classes shows slightly different survival rates, with Myo/Luminal A having the best prognosis; better than PAM50 Luminal A. TP63 expression (a known myoepithelial marker; see Figure 6) somewhat robustly defines the Myo/Luminal class. Kaplan-Meier survival analysis plots are shown comparing the survival probabilities between TP63+ and TP63-phenotypes across the whole METABRIC cohort. TP63+ confers a survival advantage comparable to that of PGR+.

**Figure 6:**
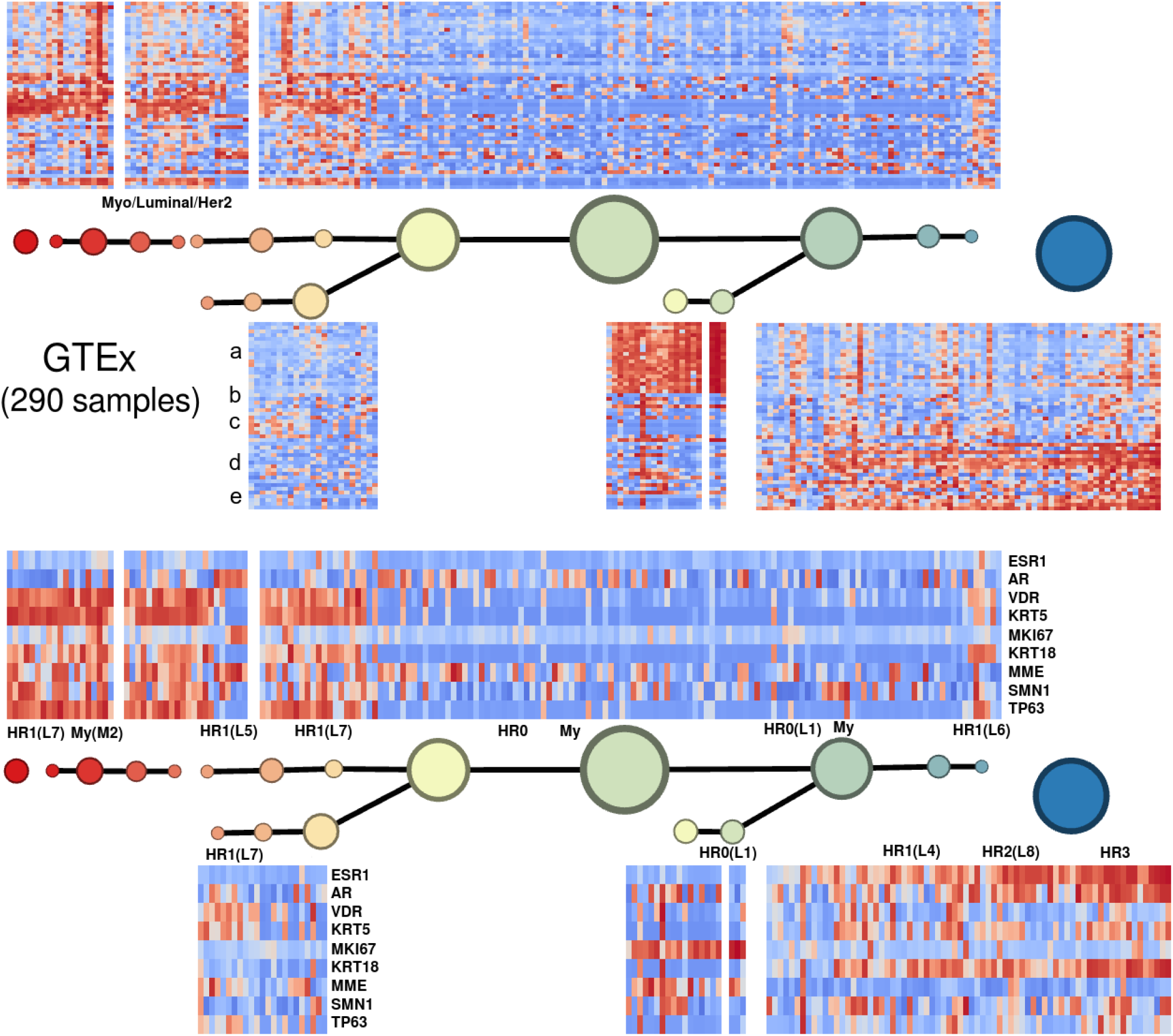
(Above) The Mapper analysis of the 290 GTEx normal mammary tissue samples, along the PAM50 gene set, using the basal-luminal score as filter function. (Below) The same sample set, in the same order, showing the expression of the marker genes of Santagata *et al*. [5] which define the normal mammary cell type classification proposed by those authors. A substantial group displays the Myo/Luminal/Her2 phenotype. According to the Santagata *et al*. classification, these samples are primarily a combination of the myoepithelial type M2 (TP63+/KRT5+) and the luminal-epithelial type L7 (VDR+/KRT5+).

The Myo/Luminal and Myo/Luminal/Her2 subtypes have signatures with the most new features. Kaplan-Meier analysis shows that the Myo/Luminal A (FOXC1-/MIA-/PHGDH-) phenotype has the best prognosis of all, with 93% survival at 5 years (Figure 5).

To investigate the Myo/Luminal class further, we drew upon the classification of normal mammary cell types of Santagata *et al*. [5] in terms of marker genes/proteins ESR1, AR, VDR, KRT5, MKI67, KRT18, MME, SMN1, and TP63. Figure 6 shows the Mapper analysis of the 290 normal breast tissue samples of the GTEx RNA expression database [12]. We found normal tissue expression patterns were similar to one of our class’ signatures along the PAM50 and also similar to one of the cell type patterns of Santagata *et al*. [5] along their marker genes. One of the clearest patterns was activation of only the basal gene group along the normal cell type denoted L1, characterized by expression of the proliferation marker MKI67. In addition, a clear subset of samples, displaying a superposition of the pattern of normal myoepithelial cell type M2 and normal cell type L7 (KRT5+/VDR+), also displayed the signature Myo/Luminal/Her2. The main characteristic of M2 is expression of TP63. We found that TP63 expression can be used as a single marker for the Myo/Luminal class (specificity: 80.8%, sensitivity: 82.8%).

### Basal/Myoepithelial (triple-negative) subclass with immune-related survival advantage (Figure 7)

Since the Myo/Luminal class is heterogeneous with respect to FOXC1, MIA, and PHGDH expression, we expected that FOXC1+/MIA+/PHGDH+ would be associated with a more aggressive phenotype. After all, these genes are highly expressed in the PAM50 Basal subtype (Basal/Myo). We found that while this is true for the first 48 months after diagnosis, the FOXC1+, MIA+, and PHGDH+ phenotypes all showed very favorable survival rates *contingent on survival to 48 months* (Figure 7). We hypothesized that this phenomenon might generalize to the PAM50 Basal subtype. To test this, we sought genes from the set of 18,543 genes available for the METABRIC cohort which would separate the long-term and short-term survivors in the FOXC1+/MIA+/PHGDH+ group. The 100 most significant genes with respect to the t-test for difference of mean expression (−log_10_(*p*) value greater than 6.7) included the genes coding for the B-cell antigen receptor complex-associated protein alpha and beta chains, the B-cell-specific coactivator OBF-1, the pre-B lymphocyte-specific protein-2, and B-cell maturation factor (CD79A, CD79B, POU2AF1, IGLL1, and TNFRSF17), as well as CD38, expressed by many immune cells. (In fact, CD79A is one of the major positive expression markers for the Claudin-low subtype introduced by Prat and Perou [9]. The Claudin-low subtype and our CD79A+/CD38+/IGLL1+ type are both subgroups of the Basal group.)

**Figure 7:**
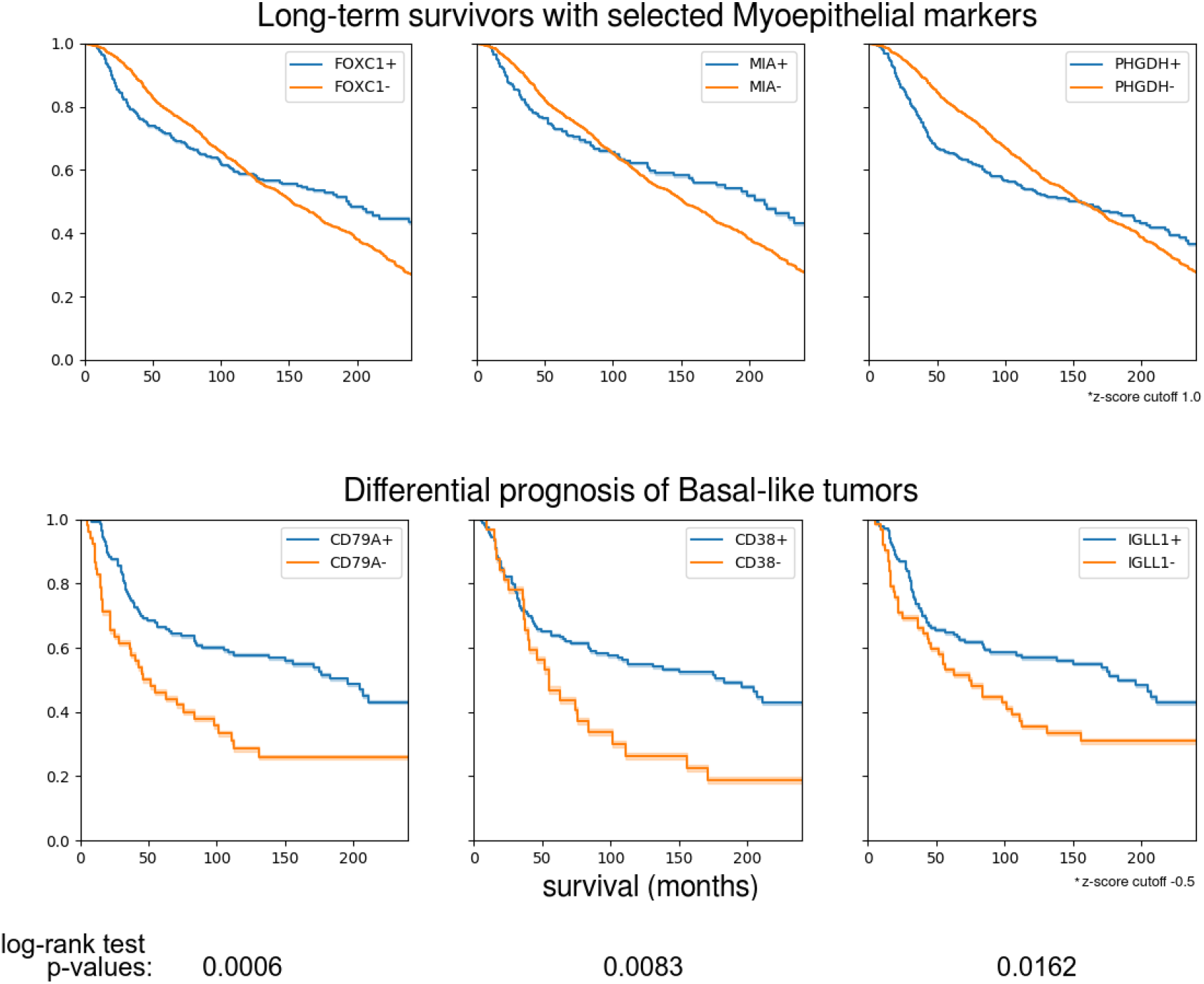
(Above) The FOXC1+/MIA+/PHGDH+ phenotype, observed in the Myo/Luminal B class but not the Myo/Luminal A class, confers a survival disadvantage for approximately the first 48 months after diagnosis, and a survival advantage afterwards. (Below) Of the top 100 genes out of 18543 exhibiting statistically significant mean differences between the FOXC1+/MIA+/PHGDH+ short-term and longterm survivors, several are B-lymphocyte-related, including: CD79A (immunoglobulin-alpha), CD38, and IGLL1 (immunoglobulin lambda-like polypeptide 1). FOXC1+/MIA+/PHGDH+ is also observed in the PAM50 Basal subtype. Within the Basal subtype, CD79A+, CD38+, and IGLL1+ confer a significant survival advantage after 48 months.

Figure 7 shows that expression of each of CD79A, CD38, and IGLL1 strongly stratifies the Basal tumors into a poor prognosis group and another group with much better prognosis after 48 months.

## 3 Discussion

### Clearly-defined 50-gene signatures

Only certain combinations of the elementary phenotypes we identified, Basal, Luminal, Myoepithelial, and Her2 are observed in breast tumors. For example, the Luminal/Basal, Basal/Myo, and Myo/Luminal are all observed, but the combination Luminal/Basal/Myo is not. We conclude that in the tumor development process, the activation of any two of the Luminal, Basal, and Myoepithelial gene groups precludes the further activation of the third.

### Myo/Luminal class with good survival

Some of the genes in the new Myoepithelial gene group (denoted b in Figure 2) concurrently stratify the Myo/Lum class. These include FOXC1, MIA, and PHGDH. The protein product of PHGDH, phosphoglycerate dehydrogenase, is a key enzyme participating in biosynthesis of serine. Maddocks *et al*. [13] find that functioning p53 is required for complete activation of the serine synthesis pathway in human cancer cells. Since the Myo/Luminal tumors have a very low TP53 mutant rate of only 15.6% in comparison to 78% for Basal/Myo, the Myo/Luminal tumors, with functioning p53, are probably capable of synthesis of serine in response to serine starvation. Only the Myo/Luminal B subclass of Myo/Luminal actually expresses PHGDH, suggesting serine synthesis and metabolism. Since Myo/Luminal A exhibits better survival rates than Myo/Luminal B, our findings are consistent with the results of Labuschagne *et al*. [14] and Amelio *et al*. [15] implicating serine metabolism in promoting tumor growth.

TP63 is one of the myoepithelial markers in the work of Santagata et *al*. [5] and also a key marker for our Myo/Luminal class. From the Kaplan-Meier analysis in Figure 5 we conclude that TP63 expression confers a survival advantage even greater than the well-known survival advantage conferred by PGR expression across the whole METABRIC cohort.

The PAM50 subtype most resembling the Myo/Luminal class is Normal-like. The status of the Normal-like subtype has been uncertain since its introduction by Perou *et al*. [4]. It is often thought to represent non-cancer tissue which is incidentally present in bulk tissue samples. For example, the PAM50 classifier uses actual normal tissue samples to train the centroid of the Normal class. However, in our analysis all of the classes of breast cancer show similarity to some combination of normal mammary cell types.

### Basal/Myoepithelial (triple-negative) subclass with immune-related survival advantage

We found a significantly lower death rate after 4 years for patients with basal tumors expressing key B-lymphocyte related markers CD79A, CD79B, POU2AF1, IGLL1, and TNFRSF17. This group is 80.3% of all patients with basal tumors surviving to 4 years. We conclude that the remaining 19.7% of these patients, with basal tumors lacking these markers, are still at high risk of mortality. This observation is consistent with the finding of Rueda *et al*. [16] that a certain subgroup of triple-negative breast cancers can be defined which rarely recurs after 5 years.

## 4 Future work

Responses to specific drugs or therapies should be investigated to decide whether some patients with Luminal but not Myo/Luminal tumors are undertreated.

Moreover, future work should address the question of why the 4 main gene groups appear. One possible explanation is that the 4 prototypical expression patterns Luminal, Basal, Myoepithelial, and Her2-related represent types of clones derived from an original transformation, and the combinations of these prototypes correspond to a certain clonal mixture. Another possibility is that the observed expression patterns are superpositions of actual tumor expression, expression of tumor microenvironmental normal cells with types related to the 4 prototypes, or expression patterns similar to original normal ancestor cells. New techniques of single-cell sequencing, potentially in conjunction with tumor-level spatial mapping, may provide answers to these questions.

Finally, the differential prognosis among triple-negative tumors observed with respect to the B-lymphocyte-related stratification suggests that the immune systems of approximately 51% of patients with triple-negative tumors can naturally and reliably mount a successful response to the tumor. If this hypothesis is correct, a longitudinal study monitoring the immune system of triple-negative patients should be able to discover exactly what response is mounted, which could lead to a new therapy which induces this natural response.

## 5 Methods

Topological Data Analysis (TDA) methods, employing ideas from the mathematical field of topology, have gained popularity in recent years. More precisely, discrete algorithmic counterparts of topological concepts have emerged in response to the availability of large datasets harboring hidden structures. Mapper [17], a discrete analogue of a Morse-theoretic analysis of a manifold with respect to a height function, or “filter” function, has received particular attention with regards to both its theoretical foundations [18,19] and, following Nicolau *et al*. [10], its application to cancer genomics [20–22]. Mapper builds a graphical summary of a given sample set with respect to a chosen stratification (filter) function. See the *Supplementary Information* for a detailed description of our Mapper analysis method.

We use three sample sets: TCGA, METABRIC [23,24], and GTEx [12]. The 1082 TCGA and 1904 METABRIC mRNA expression *z*-score data sets along the PAM50 gene set were retrieved from cBioPortal [25,26]. The 290 GTEx normal breast data set was downloaded from the GTEx portal.

The “filter function” or initial stratification is taken to be a basal-luminal epithelial differentiation score, calculated as the average expression z-score of luminal-epithelial markers (XBP1, FOXA1, GATA3, ESR1, ANXA9) minus the average expression z-score of basal-epithelial markers (KRT17, KRT5, DST, ITGB4, LAMC2, CDH3, LAD1, ITGA7). Selected largely on the basis of Perou *et al*. [4], the basal markers are all associated with anchorage of epithelial cell layers to the basement membrane, while the luminal markers are all expressed in well-differentiated or mature luminal epithelial cells.

The Mapper graph and 50-gene signatures determined from the METABRIC breast tumor samples are shown in Figure 3. Correlation-based clustering along small contiguous subsets with respect to the graph yielded the 5 main gene groups.

A simple classifier was constructed from the table of observed signatures (see Figure 3) as follows: For a given sample and a given signature or profile, the average values for each gene group are calculated, then added together with the signature signs as weights. The resulting number is a similarity score between the sample and the signature. The sample is assigned to the highest-scoring signature.

Finally, the classes and gene groups shown in Figure 2 were adjusted: The two myoepithelial gene groups were merged, the Myo/Luminal A and Myo/Luminal B classes were merged as a result, and Luminal expression was used to delineate classes Basal/Her2 and Basal/Luminal/Her2.

## Data and code availability

All the data used in this study is publicly available. The code is available upon request.

## Contributions

JCM performed research and analyzed data. JCM and SN drafted the manuscript. AL contributed a number of key suggestions for the present version. SN, AL, MP, JOD, and AT edited the paper. AT and JOD directed the research.

## References

[1] Duffy, M. et al. Clinical use of biomarkers in breast cancer: Updated guidelines from the european group on tumor markers (egtm). European journal of cancer 75, 284–298 (2017).

[2] Coates, A. S. et al. Tailoring therapies-improving the management of early breast cancer: St Gallen International Expert Consensus on the Primary Therapy of Early Breast Cancer 2015. Ann. Oncol. 26, 1533–1546 (2015).

[3] Untch, M. et al. Primary therapy of patients with early breast cancer: Evidence, controversies, consensus. Geburtshilfe und Frauenheilkunde 75, 556–565 (2015).

[4] Perou, C. M. et al. Molecular portraits of human breast tumours. Nature 406, 747–752 (2000).

[5] Santagata, S. et al. Taxonomy of breast cancer based on normal cell phenotype predicts outcome. J. Clin. Invest. 124, 859–870 (2014).

[6] Gudjonsson, T., Adriance, M. C., Sternlicht, M. D., Petersen, O. W. & Bissell, M. J. Myoepithelial cells: their origin and function in breast morphogenesis and neoplasia. J Mammary Gland Biol Neoplasia 10, 261–272 (2005).

[7] Sorlie, T. et al. Gene expression patterns of breast carcinomas distinguish tumor subclasses with clinical implications. Proc. Natl. Acad. Sci. U.S.A. 98, 10869–10874 (2001).

[8] Parker, J. S. et al. Supervised risk predictor of breast cancer based on intrinsic subtypes. J. Clin. Oncol. 27, 1160–1167 (2009).

[9] Prat, A. & Perou, C. M. Deconstructing the molecular portraits of breast cancer. Mol Oncol 5, 5–23 (2011).

[10] Nicolau, M., Levine, A. J. & Carlsson, G. Topology based data analysis identifies a subgroup of breast cancers with a unique mutational profile and excellent survival. Proc. Natl. Acad. Sci. U.S.A. 108, 7265–7270 (2011).

[11] Rousseeuw, P. J. Silhouettes: a graphical aid to the interpretation and validation of cluster analysis. Journal of computational and applied mathematics 20, 53–65 (1987).

[12] Lonsdale, J. et al. The Genotype-Tissue Expression (GTEx) project. Nat. Genet. 45, 580–585 (2013).

[13] Maddocks, O. D. et al. Serine starvation induces stress and p53-dependent metabolic remodelling in cancer cells. Nature 493, 542–546 (2013).

[14] Labuschagne, C. F., van den Broek, N. J., Mackay, G. M., Vousden, K. H. & Maddocks, O. D. Serine, but not glycine, supports one-carbon metabolism and proliferation of cancer cells. Cell Rep 7, 1248–1258 (2014).

[15] Amelio, I., Cutruzzola, F., Antonov, A., Agostini, M. & Melino, G. Serine and glycine metabolism in cancer. Trends Biochem. Sci. 39, 191–198 (2014).

[16] Rueda, O. M. et al. Dynamics of breast-cancer relapse reveal late-recurring er-positive genomic subgroups. Nature (2019). URL https://doi.org/10.1038/s41586-019-1007-8.

[17] Singh, G., Memoli, F. & Carlsson, G. Topological Methods for the Analysis of High Dimensional Data Sets and 3D Object Recognition. In Botsch, M., Pajarola, R., Chen, B. & Zwicker, M. (eds.) Eurographics Symposium on Point-Based Graphics (The Eurographics Association, 2007).

[18] Carrierè, M. & Oudot, S. Structure and stability of the one-dimensional mapper. Foundations of Computational Mathematics (2017). URL https://doi.org/10.1007/s10208-017-9370-z.

[19] Dey, T. K., Mmoli, F. & Wang, Y. Multiscale Mapper: Topological Summarization via Codomain Covers. In Proceedings of the Twenty-Seventh Annual ACM-SIAM Symposium on Discrete Algorithms (2016).

[20] Lum, P. Y. et al. Extracting insights from the shape of complex data using topology. Sci Rep 3, 1236 (2013).

[21] Lockwood, S. & Krishnamoorthy, B. Topological features in cancer gene expression data. http://arxiv.org/abs/1410.3198v1 (2014).

[22] Jeitziner, R. et al. Two-tier mapper: a user-independent clustering method for global gene expression analysis based on topology. https://arxiv.org/pdf/1801.01841.pdf (2017).

[23] Curtis, C. et al. The genomic and transcriptomic architecture of 2,000 breast tumours reveals novel subgroups. Nature 486, 346–352 (2012).

[24] Pereira, B. et al. The somatic mutation profiles of 2,433 breast cancers refines their genomic and transcriptomic landscapes. Nat Commun 7, 11479 (2016).

[25] Gao, J. et al. Integrative analysis of complex cancer genomics and clinical profiles using the cBioPortal. Sci Signal 6, pl1 (2013).

[26] Cerami, E. et al. The cBio cancer genomics portal: an open platform for exploring multidimensional cancer genomics data. Cancer Discov 2, 401–404 (2012).

