## Supplementary Information for "Robust and Interpretable PAM50 Reclassification Exhibits Survival Advantage for Myoepithelial and Immune Phenotypes"

### Supplementary Information: Mapper-based unsupervised analysis of high-dimensional point clouds

This supplementary document is a detailed description of the workflow we used to calculate Mapper-based graphical summaries of high-dimensional point clouds and generate heatmap-enriched visual representations of the resulting graphs. Though we use the language of RNA expression profiles, the method is agnostic with respect to the interpretation of the numerical quantities of the input data sets.

#### 1 Overview

A chosen filter function provides a rough stratification of a given sample set. Each stratum is clustered into nodes with edges between nodes representing cluster overlaps. The choice of filter function is important because it provides the overall basis for dimensional reduction to a 1D network or graph, and hence the basis for interpretation of the results with respect to spatial arrangement in the graph. Possible choices include generic functions like centroid-distance or eigenfunctions of a distance-weighted-graph Laplacian, but also quantitative outcomes or covariates of interest like disease grade. In our work, the filter function is informed by biologically relevant features.

The range of filter values is covered by overlapping bins. Though more sophisticated covers may be chosen, as in Dey et al. [?], we employ a simple uniform data-driven cover for a given number of bins and a given percentage of overlap between successive bins. This percentage is defined as the ratio of the number of samples in the overlap to the total number of samples. The number of bins is chosen approximately equal to the square root of the total number of samples, in order that there are approximately as many samples in each bin as there are bins. This tethers the two quantities together, ensuring that in general the number of bins and the bin sizes themselves are neither too large nor too small. The percentage, which roughly controls the sparsity of the resulting graph, should be chosen between the levels which fully connect and fully disconnect the graph.

Next the samples in each bin are clustered according to an established clustering method. In the present work, Ward's method [?] is employed, approximately minimizing within-cluster variances. The clusters, or nodes, are joined by edges whenever they overlap by some subset of samples. The resulting graph can be given a planar representation using graph visualization software systems like Gephi or Cytoscape. We arrange nodes so that the horizontal axis reflects the values of the filter function. For the Mapper algorithm, we used the Müllner implementation Python Mapper.

Heatmap representations of the original data values for each sample are overlaid on the graphical diagram, where each sample's representation is placed near the location of that sample along the graph. For interpretability, an ordering of the coordinates (genes) should be selected carefully. Convenient orderings are found by grouping together genes which are coexpressed in several nodes.

### 2 Detailed pipeline

The workflow requires:

1. Python 3, with several additional libraries, including: matplotlib, numpy, and scipy.
2. Gephi (graph analysis and visualization software).
3. A modified version of Daniel Müllner's Python Mapper.

Steps:

1. Select a gene set. One large database of gene sets is MSigDB.
2. Obtain a CSV file of sample values for exactly that gene set (samples as rows, genes as columns; with row and column names). A convenient source for RNA expression and methylation profiles, among other data, is cBioPortal. Note that cBioPortal alphabetizes the gene names. Name the CSV file to reflect the idiosyncrasies of the source, for reproducibility. It is convenient to include the number of samples in the filename, so that if missing values force omission of some samples, this fact is reflected in filename differences.
3. (Optional) Obtain another CSV file with the same exact sample set, in order, but with different data or genes (if needed for an auxiliary analysis, e.g. the calculation of the filter function).
4. Make a raw version of the main chosen CSV file. (With no header or sample names. This is required by Mapper.)
5. Calculate a filter function, one numerical value for each sample, and save it as a CSV file. This is entirely application-specific. Make two versions: One with sample names as the first column, and one raw version without this column.
6. Run modified Mapper, selecting the raw expression data set and the raw filter function file.
7. Select the parameters. Choose the number of bins to be approximately the square root of the number of samples. Choose the "gap size" to be between approximately 0.1 and 0.4. $<0.1$ tends to force each sample of a given bin to reside in its own

cluster, while >0.4 tends to put all samples of a given bin into the same cluster. Choose the overlap percentage between about 20% and 50%. >50% tends to create a dense connection network, in addition to the possibility of a large number of higher-dimensional simplices in the resulting simplicial complex. There is no standardized interpretation methodology for such complexes, so use less than 50% to make sure to get a sparse graph. <20% tends to create a network which is too sparse. For clustering method, use Ward clustering.

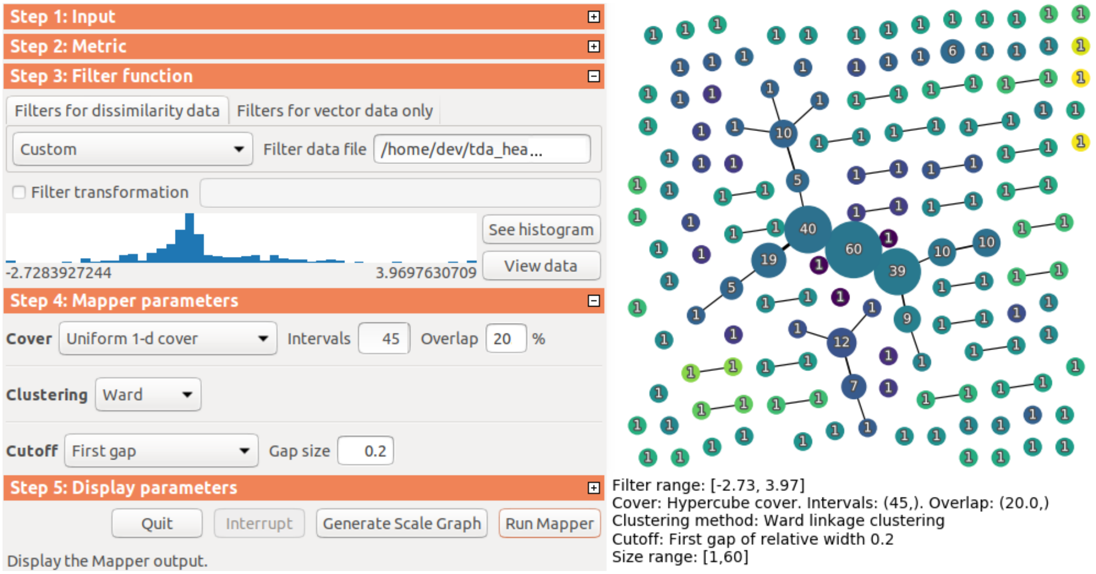


8. Save an image file with the Mapper-displayed graph. On the image, choose and mark out regions of interest (with labels, e.g. A, B, ...). We use a decomposition of the graph into long filter-directed paths, starting with the path with the largest total number of samples (the sum of the number of samples in each node along the path).
9. Save the integer indices of the sample set of each region of interest, as in `A.txt`, `B.txt`, ..., by mouse selection (Shift-click for multiple nodes) in the Mapper viewing window and then key combination Ctrl-Shift `s`.
10. Order the samples by membership in the marked sample sets, and then by filter function value.
11. Use matplotlib in Python to render heatmaps of the original expression values, grouped by the marked sample sets and ordered by filter value.
12. Import the Mapper graph into Gephi. We have modified Mapper to export the graph information to files with the name format `nodes...csv` and `edges...csv`. Choose *Import Spreadsheet* to import the nodes file, then choose *Import Spreadsheet* again to import the edges file.
13. Find a convenient display for the graph using the editing tools. Use the "force atlas" as a preliminary layout, make the filter function apparent as the left-to-right dimension,

make node size correspond to the number of samples in the corresponding sample set, remove extraneous boundary effects like singleton nodes, etc. Export the graph view image.

14. Label the graph image with the rendered heatmaps.

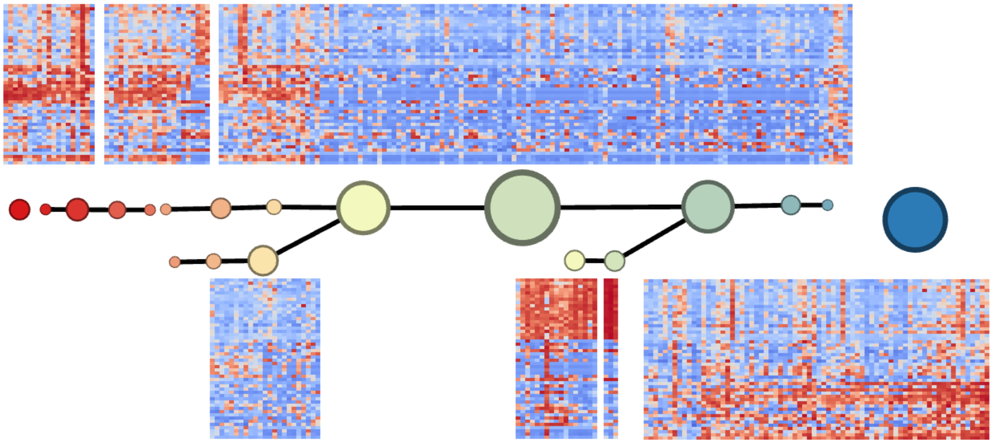

#### References


[1] Tamal K. Dey, Facundo Mmoli, and Yusu Wang. Multiscale Mapper: Topological Summarization via Codomain Covers. In *Proceedings of the Twenty-Seventh Annual ACM-SIAM Symposium on Discrete Algorithms*, 2016.

[2] Joe H. Ward, Jr. Hierarchical grouping to optimize an objective function. *J. Amer. Statist. Assoc.*, 58:236–244, 1963.
